# Epigenetic Crosstalk Between BCG-Infected Macrophages and Naïve Monocytes Potentiates Antimycobacterial Activity

**DOI:** 10.64898/2026.01.12.698601

**Authors:** Lovisa Ellinger, Shamila D. Alipoor, Shumaila Sayyab, Sadaf Kalsum, Maria Lerm

**Author notes:** Lovisa Ellinger and Shamila D. Alipoor contributed equally to this work. **Corresponding author** Maria Lerm, Div. of Inflammation and Infection, Lab 1, floor 12, Dept. of Biomedical and Clinical Sciences, Faculty of Medicine and Health Sciences, Linköping University, SE-58185 Linköping, Sweden.

## Abstract

Tuberculosis (TB), caused by *Mycobacterium tuberculosis* (Mtb), remains a leading cause of death from infectious disease. Infected cells secrete extracellular vesicles (EVs), nanosized membrane-bound particles containing bioactive molecules known to mediate intercellular communication and influence immune regulation during infection. We hypothesized that EVs secreted from *Mycobacterium bovis* Bacillus Calmette Guérin (BCG)-infected macrophages could epigenetically reprogram naïve monocytes and enhance their mycobactericidal activity against Mtb. A transwell co-culture system was used to enable communication via EVs and soluble factors, between BCG-infected macrophages and naïve monocytes (recipient macrophages). To assess the impact of these factors, we examined epigenetic reprogramming and the ability to control Mtb growth. BCG-infected macrophages released EVs with a distinct proteomic profile mapping to multiple tuberculosis-related pathways. Recipient macrophages exhibited altered DNA methylation patterns compared to those co-cultured with untreated, *Staphylococcus aureus*-infected or hydrogen peroxide-exposed macrophages. The proteomic cargo of EVs and differentially methylated genes in recipient cells showed significant interactions with TNF as a central hub enriched in both the phagosome and tuberculosis pathway. The epigenetic reprogramming was accompanied by a trend towards improved control of Mtb *in vitro*. BCG infection induces release of EVs that, together with soluble factors, induce targeted epigenetic remodeling in recipient macrophages.

In this study, we demonstrate that extracellular vesicles (EVs) released during *Mycobacterium bovis*-infection carry a directed proteomic cargo mapping to several tuberculosis-related pathways. Furthermore, we show that these proteins interact with differentially methylated genes in EV-recipient macrophages. Our findings indicate that infected macrophages transmit epigenetic information via EVs. This work provides new insights into intercellular and epigenetic mechanisms linked to innate immune memory.

## Introduction

Tuberculosis (TB), caused by *Mycobacterium tuberculosis* (Mtb), has remained the leading cause of death from a single infectious agent for most of the past decade. Its global mortality burden is of the same order of magnitude as that of COVID-19 during the peak years of the pandemic (2020–2022), when COVID-19 temporarily surpassed TB as the foremost cause of infectious mortality. Tuberculosis primarily affects the lungs and is transmitted via aerosols released when infected individuals cough or sneeze. The immune responses to Mtb are highly heterogenous; although approximately 25% of the global population is estimated to be infected, the majority will never develop disease [1]. TB represents a spectrum ranging from exposure and bacterial clearance to latent infection, subclinical disease, and ultimately active TB [2]. Recent studies suggest that part of this heterogeneity may be attributed to differences in trained immunity. Trained immunity is a form of innate immune memory where cells are epigenetically reprogrammed after an initial microbial exposure, such as the Bacillus Calmette-Guérin (BCG) vaccination, resulting in a long-lasting enhanced capacity to respond to subsequent infections [3]. How these signals are transmitted between cells is unclear, but one potential mediator is extracellular vesicles (EVs). EVs are small lipid bilayer particles that are released by nearly all cell types and play a crucial role in intercellular signalling by transferring nucleic acids, proteins and lipids between cells [4]. They are secreted both under physiological and stressed conditions, and their molecular cargo reflects the state of the sender cell and are capable of inducing specific phenotypes in recipient cells [5, 6]. EVs can be categorized into exosomes, microvesicles, and apoptotic bodies based on their biogenesis [7]. However, the origin of isolated EVs is hard to determine and the most recent MISEV2023 guidelines suggests two new categorize based on size instead of biogenesis. Small EVs are <200 nm and large EVs are >200 nm [7]. EVs are increasingly recognized as potential mediators of epigenetic regulation through the transfer of DNA methyltransferase (DNMT) transcripts, long non-coding RNAs (lncRNAs), and microRNAs (miRNAs). These cargos are proposed to induce targeted, stable changes in gene expression, thereby influencing the epigenetic landscape and behaviour of the recipient cell [8]. Several studies have linked EV-associated mRNA, miRNA and lncRNA to epigenetic modulation processes, reviewed in [9].

Beyond their proposed role in epigenetic modulation, EVs have important immunoregulatory functions. Subsequent studies revealed that EVs contribute to antigen presentation through several mechanisms, including cross-dressing, whereby EVs are taken up by or attached to dendritic cells (DCs), enabling the display of transferred human leukocyte antigens (HLA) class II molecules on their surface [10, 11]. This mechanism is particularly important in contexts where macrophages (MΦ) or DCs are infected with intracellular pathogens, such as Mtb, and exhibit a reduced capacity for canonical antigen presentation [12]. Beyond antigen presentation, EVs also modulate innate immune responses during host-pathogen interactions. MΦ infected with intracellular pathogens, such as Mtb, *Mycobacterium bovis* (Bacillus Calmette-Guerin BCG) or *Salmonella typhimurium,* release EVs containing pathogen-associated molecular patterns (PAMPs), which activate toll-like receptors (TLRs) and myeloid differentiation factor 88 (MyD88) signalling in recipient cells [13]. Through this mechanism, EVs propagate immune signalling to distal sites, activating uninfected immune cells [14]. EVs derived from Mtb- or BCG-infected MΦ (BCG-I MΦ) carry mycobacterial antigens such as lipoarabinomannan (LAM), early secreted antigenic target 6 kDa (ESAT-6) and antigen 85 complex (Ag85), which can stimulate MΦ, DCs and T cells *in vivo* [13, 15]. These vesicles also promote the production of pro-inflammatory cytokines and induction of autophagy, key components of host defence [16–18]. However, Mtb is known to evade host immunity by subverting immune regulatory pathways to prolong its intracellular survival. EV secretion appears to be part of this strategy, EVs from Mtb-infected MΦ have been shown to suppress interferon-γ (IFN-γ) production and promote MΦ polarization towards anti-inflammatory phenotypes in distal cells [19, 20].

In this study, we investigated the epigenetic basis of stable phenotypic reprogramming in naïve monocytes. We hypothesized that EVs from BCG-I MΦ induce epigenetic changes that alter recipient cell functions. To model this interaction, BCG-I MΦ were co-cultured with naïve monocytes during differentiation to MΦ (recipient MΦ) allowing communication via soluble factors. Functional and epigenetic outcomes were then assessed. *Staphylococcus aureus* (SA) was included as an infectious control, and hydrogen peroxide (H_2_O_2_) served as a non-infectious stressor to determine the BCG-specificity of the observed effects.

## Results

### Isolation and characterization of macrophage-released EVs

EVs released by primary human MΦ infected with BCG, SA, or exposed to H_2_O_2_ were isolated from culture supernatants. The EVs were characterized to confirm morphology, size, particle number, and surface markers in alignment with the MISEV2023 guidelines [7] (Supplementary Figure 1a-c). We observed that BCG-I MΦ displayed enhanced release of EVs, accompanied by an altered surface marker expression (Figure 1a-b). Nanoparticle tracking analysis (NTA) revealed a significant increase of EVs in all treatments as compared to untreated cells with the strongest enhancement in BCG-I MΦ (Figure 1a) (One-way ANOVA and Dunnett’s multiple comparisons test (* *P* < 0.05, ** *P* < 0.01, *** *P* < 0.001)). The expression of 38 EV surface markers was analyzed, showing treatment-dependent differences between bacterial infections (BCG, SA) and oxidative stress by H_2_O_2_ (Supplementary Figure 2). Markers related to innate immune activation and antigen presentation, including CD11c, CD14, CD29, CD40, CD45, CD49e, CD86, and HLA class I, showed significantly higher mean fold changes in BCG- and SA-infected compared to H_2_O_2_-exposed MΦ, reflecting a distinct EV phenotype under oxidative stress rather than an active infection (Figure 1b) (One-way ANOVA and Dunnett’s multiple comparisons test (* *P* < 0.05, ** *P* < 0.01, *** *P* < 0.001)).

**Figure 1.**
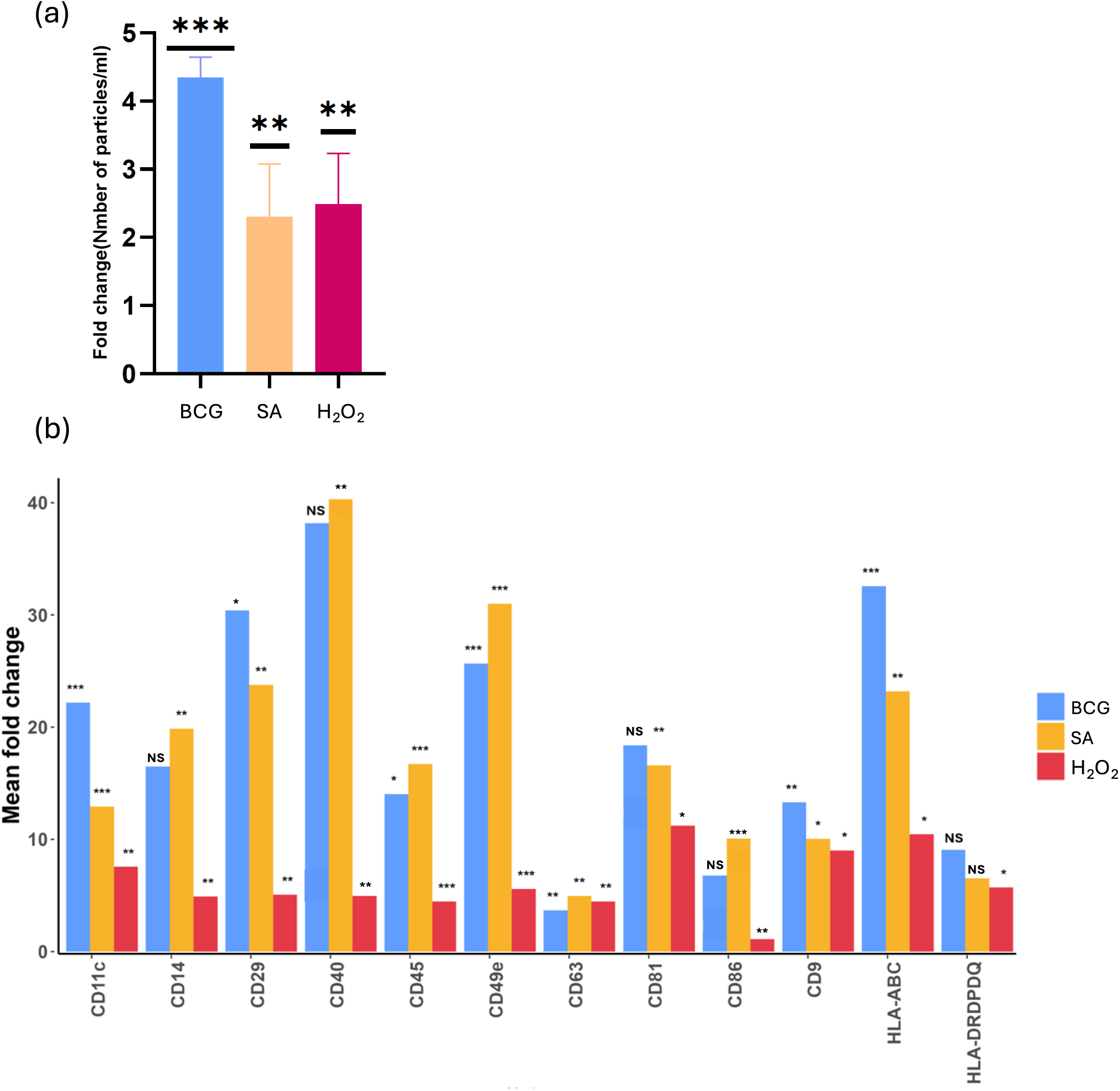
Extracellular vesicle secretion from human primary macrophages is influenced by infection with BCG or SA and exposure to H_2_O_2_. **(a)** Number of extracellular vesicle (EV) released by primary human macrophages stimulated with *Mycobacterium bovis* (BCG), Staphylococcus *aureus* (SA), hydrogen peroxide (H_2_O_2_) normalized to control. Stimulation with BCG, H_2_O_2_, and SA increased EV production compared to control, with BCG-treated cells exhibiting the highest level of EV release (One-way ANOVA and Dunnett’s multiple comparisons test (* *P* < 0.05, ** *P* < 0.01, *** *P* < 0.001)). **(b)** Surface marker expression of EVs from cells stimulated with BCG, H_2_O_2_, or SA, shown as fold change relative to control. Surface markers were detected by flow cytometry. Bar graph depicts the levels of indicated EV surface markers in the different stimulation conditions compared to unstimulated control. BCG and SA infection induce a distinct EV surface marker profile characterized by increased immune and antigen-presenting cell markers, different from oxidative stress-induced EVs in H_2_O_2_ condition. HLA-DRDPDQ is a combined bead that detects all three classical MHC class II antigens (DR, DP, DQ) and HLA-ABC targets MHC class I (A, B, C) antigens. Surface markers showing a fold change greater than 5 in at least one condition are displayed. Statistical analysis was performed using one-way ANOVA and Dunnett’s multiple comparisons test (* *P* < 0.05, ** *P* < 0.01, *** *P* < 0.001). The experiments were performed on macrophages isolated from healthy blood donors (n=10). Each donoŕs cells were treated with the respective agent (BCG, H_2_O_2_, or SA) or as control in three replicates, EVs isolated from five donors were pooled and analyzed in two consecutive experiments.

### Proteomics investigation revealed distinct protein composition for the EV released by BCG-infected macrophages

We used quantitative proteomics to identify the protein composition of the isolated EVs. A total of 935, 351, 551, and 605 peptides were identified in the EV samples from BCG-I MΦ, SA-infected-, H_2_O_2_-exposed- and untreated MΦ, respectively (Figure 2a). EVs from BCG-I MΦ the exhibited a substantial number of unique peptides, with 255 detected exclusively in this condition (Figure 2b). Gene set enrichment analysis of the genes corresponding to unique proteins identified in EVs released from BCG-I MΦ revealed an enrichment of immune-related pathways associated with intracellular infections (Figure 2c). Notably, among the top enriched were the tuberculosis pathway, phagosome maturation, endocytosis, Salmonella infection, and Leishmaniasis (Figure 2c). Network visualization of these enriched pathways demonstrated dense clusters of interconnected proteins, notably including HLA class II molecules (HLA-DQA1 and HLA-DQB1), TLR2, mannose receptor C-type 1 (MRC1) and tumor necrosis factor (TNF) (Figure 2d). In contrast, EVs from HCOC-exposed and SA-infected MΦ contained only 4 and 7 unique proteins, respectively, while 50 unique proteins were identified in EVs from untreated controls (Figure 2b). Corresponding pathway enrichment analyses revealed fewer and less pronounced enrichments for these proteins (Supplementary Figure 3).

**Figure 2.**
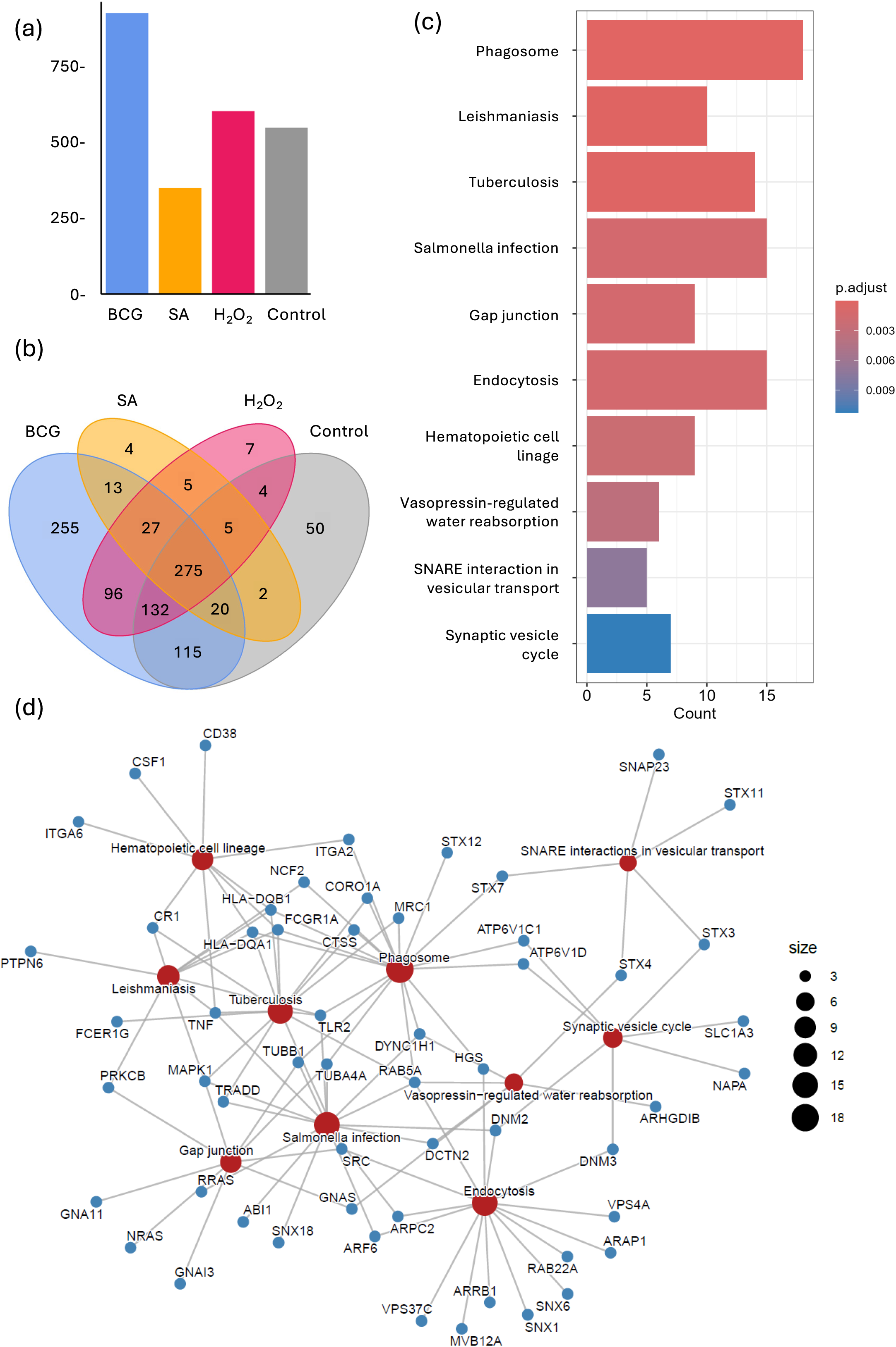
Proteomic profiles of EVs released from primary human macrophages infected with BCG, SA or exposed to H_2_O_2_. **(a)** Total number of peptides detected in each condition. Proteomic analysis was performed on extracellular vesicles (EVs) released by macrophages following stimulation with *Mycobacterium bovis* (BCG), Staphylococcus *aureus* (SA), hydrogen peroxide (H_2_O_2_), or as well as in the untreated control condition. Peptide identification was conducted using BLAST against a human cell library. The bar plot illustrates the total number of peptides identified in each condition. The experiments were performed on primary human macrophages isolated from healthy blood donors (n=5). Each donor cells were treated with the respective agent (BCG, H_2_O_2_, or SA) or left untreated as control. **(b)** Venn diagram of the protein overlaps among conditions. The Venn diagram shows the number of peptides shared or unique to each treatment (BCG-infection, SA-infection, H_2_O_2_-exposure or untreated controls. Each ellipse represents one condition, and overlapping regions indicate peptides shared between two or more conditions. Non-overlapping regions correspond to peptides uniquely identified in a single condition. **(c)** Bar plot showing the top enriched KEGG pathways among genes associated with peptides uniquely identified in the EVs released by BCG-infected macrophages. The top 10 pathways are presented in the graph. The x-axis indicates the number of genes associated with each pathway, and the color represents the adjusted p-value for enrichment. **(d)** Network plot illustrating the relationships between enriched pathways and their associated genes. Pathways are shown as red nodes, and genes are shown as purple nodes; edges indicate gene membership in pathways.

### Co-culture with BCG-infected macrophages induce epigenetic reprogramming in naïve monocytes

To assess whether EVs released by BCG-I MΦ can influence the epigenetic landscape of naïve monocytes, we employed a transwell co-culture system. MΦ were exposed to BCG, SA or H_2_O_2_, then washed to remove stimuli. Naïve monocytes in the lower chamber were co-cultured for seven days, allowing differentiation under the influence of soluble factors and EVs. DNA methylation profiling of these BCG-induced EV recipient MΦ revealed 1500 CpG sites linked to 253 unique differentially methylated genes (DMGs) (|logFC| > 0.1, *P* < 0.05), many involved in immune regulation (Figure 3a). Pathway enrichment analyses of these differentially methylated genes revealed enrichment in pathways related to Hippo signaling, EGFR tyrosine kinase inhibitor resistance and fatty acid biosynthesis and Th1 and Th2 cell differentiation (Figure 3b) (using nominal p-values (*P* < 0.05). Among the top 30 enriched pathways, we also identified enrichment in the tuberculosis pathway (*P* < 0.024, gene count 6) (Supplementary Table 1). Further network visualization of the top 15 enriched pathways shows several interconnected genes and directionality of the methylation pattern. Enrichment in Th1 and Th2 cell differentiation resulted from hypomethylation in the interferon-γ receptor 1 (IFNGR1) and interleukin-2 receptor α-chain (IL2RA) and hypermethylation in interleukin-12 receptor β2 (IL12RB2) and Janus kinase 1 (JAK1). Furthermore, enrichment in C-type lectin receptor signaling pathway involved in Mtb recognition was identified (Figure 3c). The enriched pathways associated with DNA methylation changes in co-cultures with SA-infected and H_2_O_2_-exposed cells are shown in Supplementary Figure 4.

**Figure 3:**
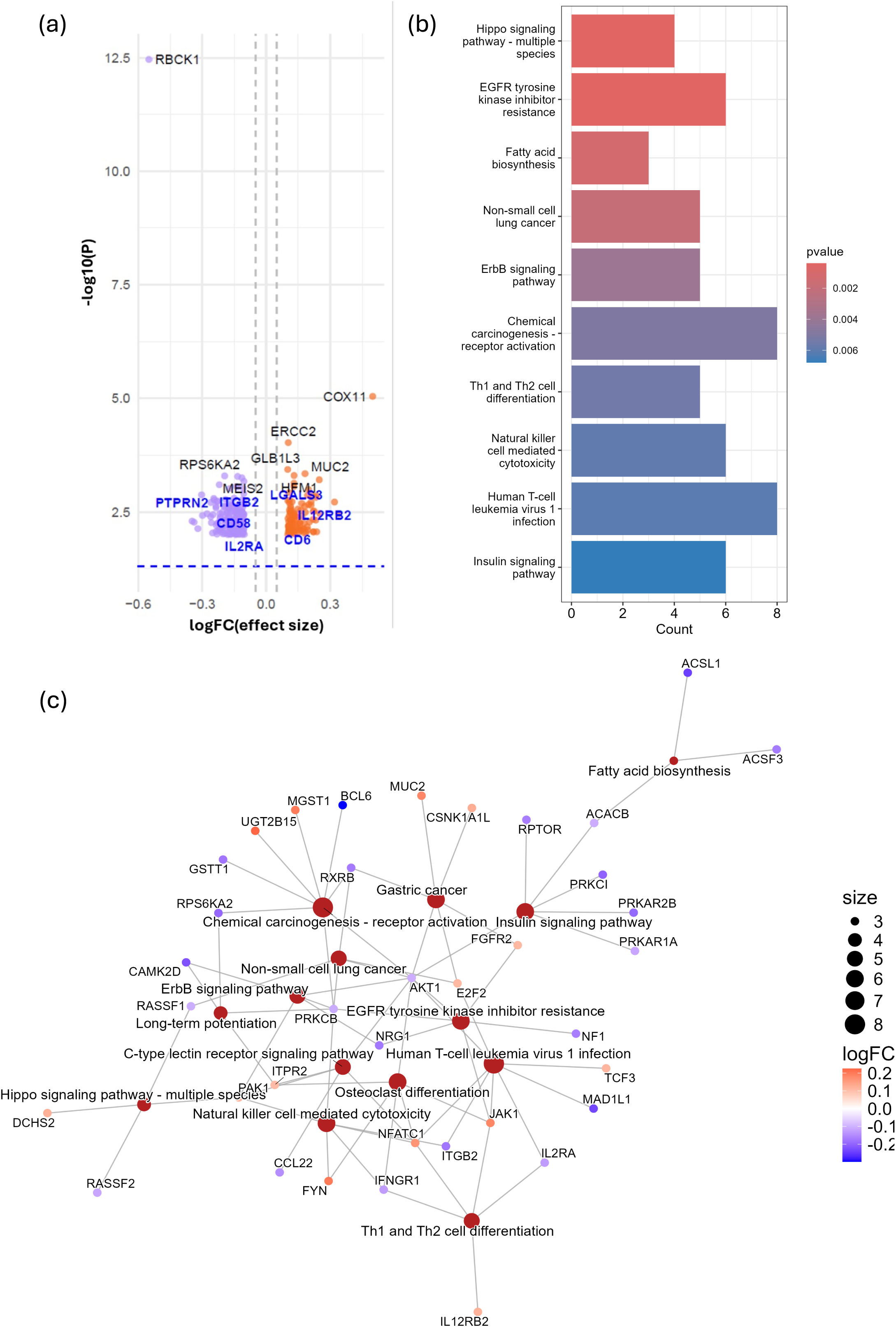
DNA methylation changes in naïve monocytes differentiated during co-cultured with BCG-infected macrophages. **(a)** Volcano plot of differentially methylated CpG sites annotated to genes showing genes (log fold change |logFC| > 0.1, *P* < 0.05). **(b)** Bar plot showing the 10 top enriched KEGG pathways among genes associated with the differentially methylated CpG sites identified in the recipient macrophages co-cultured with the *Mycobacterium bovis* (BCG)*-*infected macrophages (the top 10 pathways are shown). The x-axis indicates the number of genes associated with each pathway, and the color represents the adjusted p-value for enrichment. **(c)** Network visualization of the top 15 enriched pathways including the associated genes from DMC analysis. The network visualizes the relationships between enriched biological pathways and their associated genes. Each colored node (circle) represents either a pathway (in red) or a gene, with the gene nodes color-coded according to their logFC values. The logFC scale indicates methylation levels. Edges (lines) indicate which genes are involved in which pathways, illustrating that individual genes can contribute to multiple biological processes.

### Proteins in EVs interact with epigenetically regulated genes in recipient macrophages

To further explore the connection between the proteomic cargo of BCG-induced EVs and the epigenetic remodeling observed in the recipient MΦs, we performed a STRING analysis using a high confidence interaction score (>700). This analysis included 255 unique proteins identified in BCG-induced EVs and 253 unique DMGs (mapped to proteins) identified in the recipient MΦs, revealing a large interactome of significant protein-protein interactions (Figure 4a). TNF protein from the EVs and the DMGs CD44 and AKT1 from recipient MΦs was shown to be a central hubs with 24, 34 and 24 connections, respectively. The DMG CD44 was hypomethylated (|logFC| − 0.14) proposing increased expression of this glycoprotein in the recipient MΦ. Notably, seven proteins found in EVs (ACSL1, ASAH1, CLCN7, F13A1, PDXK, PRKAR1A, PRKCB, SLAMF8, TTYH2) were also DMGs in the recipient cells and four were present in the interactome (Figure 4a). Further KEGG pathway enrichment analysis of the high-confidence interactome showed significant overrepresentation of the phagosome, Leishmaniasis and tuberculosis pathways (Figure 4b), supporting a functional link between EV-derived proteins and epigenetic regulation in the context of intracellular infections.

**Figure 4:**
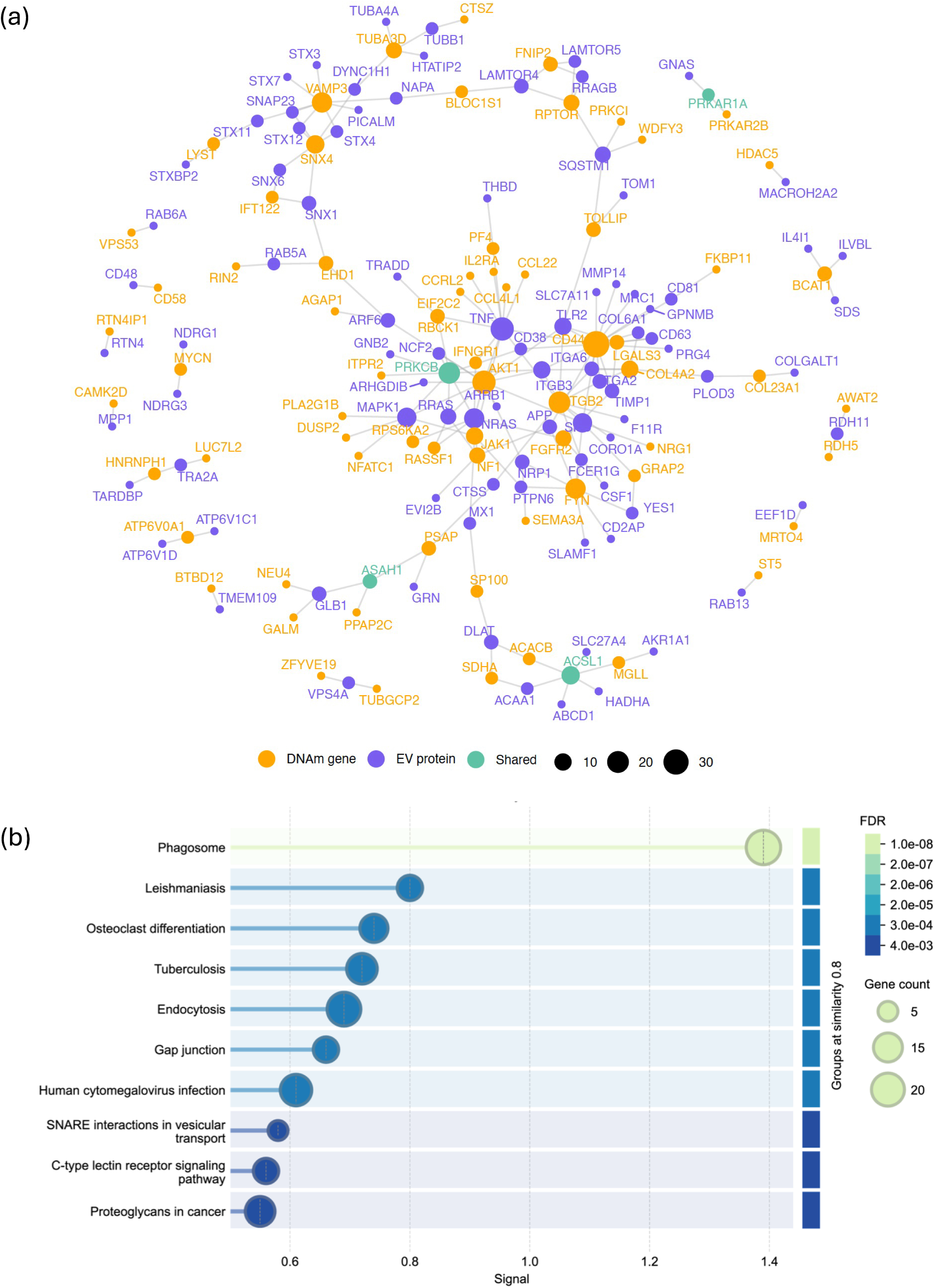
Interactome of proteomic cargo of EVs and differentially methylated genes highlighting immune regulatory hubs linked to TNF. **(a)** Protein-protein interaction network showing high-confidence (score >700) interactome of EVs protein and DNA methylation (DNAm) associated genes. Node size corresponds to degree centrality, indicating connectivity within the network. Edge transparency reflects interaction confidence. Nodes are color-coded by molecular origin: purple for EV proteins, orange for DNAm-associated genes, and green for genes shared between both datasets. **(b)** Pathway enrichment analysis of interactome. A bar plot of top 10 KEGG enriched pathways.

### Recipient macrophages exhibit a tendency of enhanced control of Mtb infection compared to controls

To evaluate whether epigenetic reprogramming translated to functional changes, we assessed the ability of recipient MΦ to control Mtb infection using a transwell system. After co-culture with BCG-I MΦ, SA-infected or H_2_O_2_-exposed MΦ, recipient MΦ were infected with GFP-expressing Mtb. Across four independent donors, recipient MΦ co-cultured with BCG-I MΦ showed a consistent trend towards reduced bacterial load over five days, with a tendency toward lower bacterial burden at day five (*P* = 0.125, Wilcoxon signed-rank test, Figure 5a-b). In contrast, cells co-cultured with SA-infected or HCOC-exposed MΦ showed variable responses across donors with most displaying increased bacterial loads at day 5 (*P* = 0.250 and 0.375, respectively; Figure 5b and growth over the five days is shown in Supplementary Figure 5). To assess whether the observed effects were mediated by EVs, EVs were isolated from the conditioned medium using ultracentrifugation, leaving other soluble factors intact. Recipient MΦ cultured in EV-depleted media showed reduced bacterial clearance compared to those exposed to EV-containing media (Wilcoxon signed-rank test, *P* = 0.125) (Figure 5c-d). In contrast, EV-removal SA- or H_2_O_2_-conditioned medium had no effect on Mtb growth (Supplementary Figure 6). These results suggest that BCG-I MΦ-released EVs contributes to anti-mycobacterial activity and is dependent on co-stimulation by other soluble factors.

**Figure 5:**
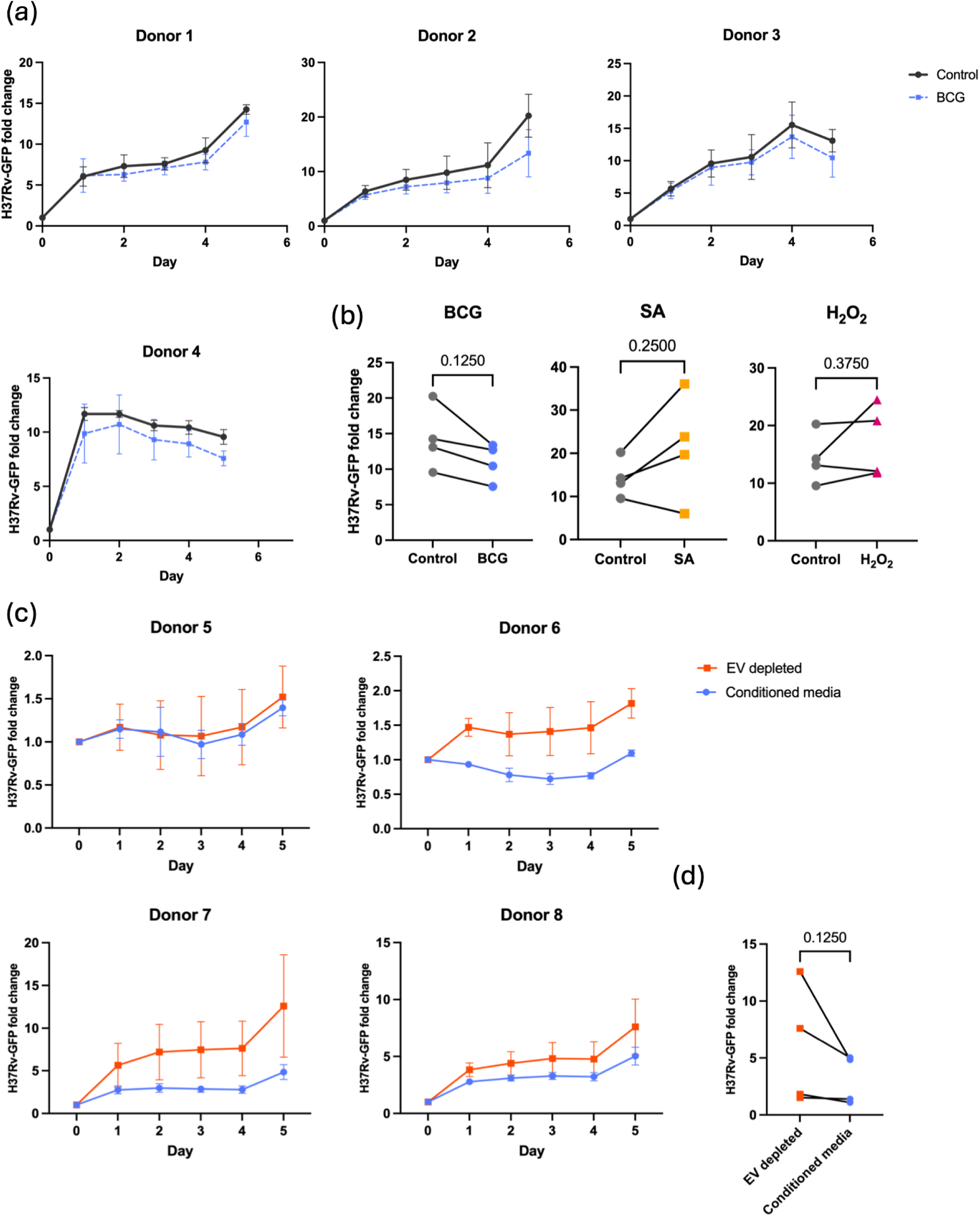
Naïve monocytes differentiated during co-culture with BCG-infected macrophages displayed an enhanced *Mycobacterium tuberculosis* clearance. **(a)** Macrophages from 4 independent donors were either untreated (control, black lines) or conditioned with EVs and soluble factors released by *Mycobacterium bovis* (BCG)-infected macrophages (BCG-I MΦ) in a transwell co-culture model (blue dashed lines) and infected with GFP-expressing *Mycobacterium tuberculosis* (Mtb). Bacterial load was monitored by live-cell imaging of GFP fluorescence over 5 days, expressed as fold change relative to the initial timepoint (day 0). **(b)** Co-culture with BCG-I MΦ lead to a tendency of decreased bacterial load at day 5 (*P* = 0.125) compared to control. Co-culture with SA infected and H_2_O_2_ exposed MΦ lead to increased Mtb load compared to controls, however, no significant differences between these conditions were detected (*P* = 0.250 and 0.375, respectively, wilcoxon signed rank test). **(c)** Monocytes from four donors were differentiated in the presence of either; ultracentrifugated EV depleted conditioned media (orange line and square) or conditioned media (blue line and dot) isolated from MΦ-cultures infected with BCG. Conditioned media contain both Evs and cytokines specifically secreted during BCG-infection. MΦ subsequently infected with GFP-expressing *M. tuberculosis* (H37Rv). Bacterial burden was assessed by quantifying the fold change in H37Rv-GFP fluorescence over five days. **(d)** At five days post-infection, MΦ cultured with EV-depleted conditioned media showed a tendency towards higher bacterial loads compared to MΦ cultured in conditioned media containing EVs (Wilcoxon signed-rank test, p = 0.125). Experiments were performed using cells from four donors, with four and three technical replicates for two donors, respectively.

## Discussion

Our study provides novel insights into the role of EVs as epigenetic modulators in the context of mycobacterial infections. We show that BCG infection induces the release of EVs with distinct surface marker profiles and a targeted proteomic cargo enriched in tuberculosis-associated proteins. When naïve monocytes were exposed to EVs and soluble factors derived from BCG-I MΦ, they underwent epigenetic remodeling in genes associated with immune activation and tuberculosis, suggesting a targeted and biologically meaningful reprogramming of recipient cells.

Proteomic analysis revealed a significant increase in EV secretion from BCG-infected macrophages compared to those exposed to *S. aureus*, H_2_O_2_, or left untreated. These EVs carried a unique set of proteins and showed marked differences in surface marker expression. Among the identified proteins were HLA class II molecules (HLA-DQA1, HLA-DQB1), several innate immune receptors including TLR2, MRC1, Fcε receptor Ig (FCER1G), Fcγ receptor Ia (FCGR1A), complement C3b/C4b receptor 1 (CR1) and TNF, molecules known to participate in pathogen recognition, antigen presentation, and inflammatory signaling. These proteins were enriched in pathways related to phagosome maturation, leishmaniasis, and tuberculosis, supporting the established role of EVs in immune activation.

Integration of EV proteomic and DNA methylation data revealed a strong interactome linking EV-derived proteins to DMGs in recipient macrophages. Several hub nodes, including TNF and CD44 occupied central positions bridging the two datasets. Immune regulatory nodes such as TNF and IFNGR1 were positioned within major hubs. Notably, the interactome was significantly enriched in the phagosome and tuberculosis pathways, underscoring the functional relevance of these molecular interactions. TNF, a key cytokine in tuberculosis, plays a central role in macrophage activation and phagosome maturation. It enhances antimicrobial activity by modulating the recruitment of Rab GTPases and other trafficking regulators, facilitating phagosome–lysosome fusion and pathogen degradation, key processes for antigen presentation and activation of T cell immunity [21, 22]. CD44 has been identified as a receptor for Mtb, along with upregulated expression in Mtb infection [23, 24]. Building on this, we observed hypomethylation of CD44 in recipient MΦ co-cultured with BCG-I MΦ, indicating increased expression of this receptor and a potential role in EV-mediated priming. These findings suggest that EVs may actively shape the functional state of recipient cells to promote effective immune responses.

These observations align with the concept of trained immunity, a hallmark of BCG vaccination, which refers to the long-term enhancement of innate immune responses through epigenetic and metabolic reprogramming [3, 25, 26]. We have previously shown that this can be induced by stimuli such as β-glucans, leading to improved control of Mtb [27]. Our current findings build on this concept by showing that EVs and soluble mediators from BCG-I MΦ, can transmit epigenetic signals to uninfected bystander cells. This supports a model in which trained immunity extends beyond the initially stimulated population via EV-mediated intercellular communication.

To explore the epigenetic landscape of recipient MΦ, we performed a DNA methylation analysis, which revealed enrichment in several immune-related and metabolic pathways. We observed enrichment in fatty acid biosynthesis, consistent with the metabolic reprogramming of activated MΦ [28]. Mtb exploits this shift by manipulating hosts lipid metabolism, promoting the formation of lipid-laden foamy MΦ, and TB patients display elevated glycerophospholipid levels [28–30]. Notably, we identified enrichment in the Th1 and Th2 cell differentiation pathway. The differentiation on naïve CD4+ T cells is crucial for mounting an efficient immune response against Mtb [31]. Interestingly, this pathway enrichment was associated with hypomethylation in the IL2RA gene, a known TNF-inducible gene that promote differentiation of T cells [32], and hypermethylation of JAK1, a kinase activated by TNF-induced phosphorylation [33, 34]. This finding suggests a mechanistic link between the TNF present in BCG-induced EVs and the epigenetically regulated immune responses in recipient MΦ. The Hippo signaling pathway, known to enhance ROS production and neutrophil recruitment [35–37], was among the top enriched. Additionally, we identified enrichment in the EGFR tyrosine kinase inhibitor resistance pathway. In infected MΦ, EGFR activation lead to STAT3 signaling which repress key antimicrobial responses including nitric oxide synthesis and the expression of pro-inflammatory cytokines such as IL-6, TNF-α, IFN-γ [38, 39]. Furthermore, genetic mutations increasing EGFR expression is associated with increased susceptibility to TB [40, 41]. Notably, we observed pronounced hypomethylation of the RBCK1 (also known as HOIL-1L) gene. Mtb protein kinase PknG has recently been shown to phosphorylate RBCK1 to block NLRP3 inflammasome assembly [42]. Together, these findings suggest that EV-mediated epigenetic remodeling affects key pathways involved in host defense, including phagosome maturation, metabolic adaptation, and cytokine regulation.

To assess the functional relevance of these epigenetic changes, we challenged recipient MΦ with Mtb. Cells exposed to soluble factors, including EVs, from BCG-I MΦ showed a consistent trend toward enhanced bacterial control. Although not statistically significant, this trend aligns with previous studies linking metabolic and epigenetic reprogramming to improved antimicrobial activity. When EVs were depleted from the conditioned media, this effect was diminished, suggesting that both EVs and other soluble factors are required for full reprogramming. This is consistent with findings by Cheng et al., who reported that EVs from Mtb-infected MΦ enhanced bacterial clearance only in the presence of IFN-γ [17]. These results support the idea that EVs act as messengers of trained immunity, priming neighboring or newly recruited monocytes for enhanced antimicrobial readiness even before direct pathogen contact. Moreover, recent studies have identified the bone marrow as a central site for the induction of trained immunity via epigenetic reprogramming of hematopoietic stem and progenitor cells (HSPCs) [43, 44]. Our results raise the possibility that EVs contribute to this systemic process by delivering immunomodulatory signals to distant compartments such as the bone marrow.

In summary, our study shows that BCG infection triggers the release of EVs with distinct surface markers and a proteomic cargo enriched in tuberculosis-associated proteins. Exposure of recipient MΦ to these EVs and soluble factors leads to epigenetic remodeling in genes related to immune activation and metabolic reprogramming. Integration of proteomic and methylation data revealed a TNF-centered interactome enriched in the phagosome and tuberculosis pathways. These findings suggest that EV uptake contributes to the establishment of trained immunity by shaping the epigenetic landscape of recipient cells.

## Material and Methods

### Study Design

This study investigated how BCG-I MΦ modulate healthy MΦ responses against Mtb, focusing on the role of EVs released by infected MΦ. A transwell co-culture model was used, where differentiated MΦ were stimulated with BCG, SA, or H_2_O_2_ on the upper chamber. Naïve monocytes in the lower chamber matured into differentiated MΦ influenced by EVs and soluble factors released by stimulated MΦ from the upper chamber. These recipient MΦ were then infected with Mtb to assess antimicrobial activity via live-cell imaging. Further analyses included profiling EVs released by infected/stimulated MΦ and epigenetic profiling of the recipient MΦ. Further methodological details can be found in the Supplementary Methods.

### Ethics Statement

Buffy coat preparations were obtained from healthy volunteers at Linköping University Hospital Blood Bank. Informed, written consent was obtained from all donors in accordance with ethical standards of the Helsinki Declaration. No ethical approval was required for this study in accordance with local and national guidelines.

### Bacterial Culture Preparation

BCG cultures were grown for 3 weeks; SA was prepared from overnight cultures with a 2-hour subculture. Bacterial suspensions were centrifuged, washed, and resuspended in antibiotic-free media. Bacterial concentrations were calculated based on optical density measurements.

### Monocyte Isolation and Macrophage Differentiation

Peripheral blood mononuclear cells (PBMCs) were isolated by density gradient centrifugation. Monocytes adhered to culture dishes and differentiated into MΦ over six days in complete DMEM supplemented with 10% FBS and M-CSF.

### Preparation of Recipient Macrophages by Co-culture

On day 6, MΦ were reseeded into transwell inserts and stimulated with BCG, SA, or H_2_O_2_. Naïve monocytes were seeded in the lower chamber, cultured together for 3 days to allow maturation.

### Control Experiments

Conditioned media were ultracentrifuged to deplete EVs. Naïve monocytes were cultured in EV-depleted or untreated media prior to Mtb infection.

### Mtb Infection and Live-Cell Microscopy

Mtb H37Rv-GFP cultured to logarithmic phase was used at MOI 1. Infected MΦ were monitored via IncuCyte live-cell imaging every 6 hours for up to 5 days, quantifying bacterial growth by fluorescence intensity.

### DNA Extraction and Methylation Profiling

DNA was extracted and DNA methylation profiling was performed using the Illumina MethylationEPIC array. Data preprocessing included normalization, filtering, batch effect correction, and identification of differentially methylated CpG sites. Pathway enrichment analyses were conducted for functional insight.

### Extracellular Vesicle Isolation and Characterization

EVs were isolated from conditioned media by size exclusion chromatography with IZON qEV columns after concentration and filtration. EV size and concentration were analyzed by NTA. Morphology was assessed by TEM, and surface markers profiled using multiplex bead-based flow cytometry.

### Proteomics of Extracellular Vesicles

Proteins were extracted from EVs, prepared by ultrasonic filter-assisted sample preparation (FASP) [45], and digested with trypsin. Peptides were analyzed via high-resolution LC-MS/MS. Protein identification utilized Spectronaut software [46]. Functional annotation and pathway enrichment were performed for proteins unique to treatment conditions.

### Statistical analysis

The number of EVs and surface marker expressions from different conditions were compared using one-way ANOVA and Dunnett’s multiple comparisons test. Statistical analyses on Mtb growth experiments were performed using Wilcoxon signed-rank test with GraphPad Prism (version 10.5.0). We used linear modeling followed by empirical Bayes moderation, as implemented in the limma package for differential analysis of methylation data. P-values were adjusted for multiple testing using the Benjamini-Hochberg procedure for False Discovery Rate (FDR) correction at 5%. In case no significant FDR was reached, we used nominal p-value < 0.05.

## Data/code Availability

The datasets generated during the current study are not publicly available due to ethical dilemmas in traceability of DNA methylation data, but processed pseudonymized data depleted of genetic variant information (beta matrixes with Illumina probe IDs and beta values) will be available upon request through the Federated European Genome-phenome Archive (FEGA) Sweden controlled-access repository, upon publication

Bioinformatic pipelines used to analyze the data and to generate graphs and figures will be available on the following https://github.com/Lerm-Lab/Epigenetic-Crosstalk

## Acknowledgements

We would like to acknowledge Clinical Genomics Linköping, Science for Life Laboratory, Linköping University for DNA methylation analysis. We would like to acknowledge the staff at Linköping Blood Bank and the anonymous blood donors.

## Author contributions

L.E, S.D.A and M.L conceptualized the study. L.E and S.A.D performed method optimizations. S.A.D and S.K performed experiments. S.A.D and L.E performed data analysis. S.D.A performed and S.S guided the bioinformatic analysis. M.L funded the study. L.E and S.D.A wrote the manuscript, and all authors contributed to the final version.

## Supplementary figure legends

**Supplementary figure 1. Extracellular vesicles (EVs) released by human primary macrophages characterized by morphology, size and surface markers. (a)** Transmission electron microscopy (TEM) image showing a spherical to cup-shaped morphology with a visible lipid bilayer membrane in all samples. **(b**) Nanoparticle Tracking Analysis (NTA) of extracellular vesicles isolated from macrophage culture supernatants, showing particle size distribution with an average size of 114 ± 3.8 nm across the samples. **(c)** Bead-based flow cytometry analysis of EVs demonstrating surface marker expression of CD9, CD63, and CD81. EVs were isolated by Izon size exclusion chromatography columns. The results shown are representative of three independent experiments conducted with macrophages derived from five different donors. Macrophages were infected with *M. bovis* (BCG), *S. aureus* (SA), exposed to hydrogen peroxide (H_2_O_2_) or left untreated.

**Supplementary figure 2. Surface marker expression of EVs from human primary macrophages infected with BCG, *S. aureus* or exposed to H_2_O_2_ shown as fold change relative to control.** Surface markers were detected by flow cytometry. Bar graph depicts the levels of indicated extracellular vesicle (EV) surface markers in the different stimulation groups compared to unstimulated control. Infection with *M. bovis* (BCG) and *S. aureus* (SA) induced a distinct EV surface marker profile characterized by increased immune and antigen-presenting cell markers, different from H_2_O_2_-induced EVs. HLA-DRDPDQ is a combined bead that detects all three classical MHC class II antigens (DR, DP, DQ) and HLA-ABC targets MHC class I (A, B, C) antigens. Experiments were performed on EVs isolated from cells of ten donors, EVs from five donors were pooled before flow cytometry analysis.

**Supplementary figure 3. Pathway enrichment of proteomic data from extracellular vesicles released by macrophages under different conditions.** KEGG pathway enrichment analysis on genes associated to peptides identified in extracellular vesicles released by human primary macrophages infected with *M. bovis* (BCG), *S. aureus* (SA), exposed to hydrogen peroxide (H_2_O_2_) or left untreated. Dot plot showing the top 5 enriched KEGG pathways for each group.

**Supplementary figure 4. DNA methylation changes in recipient macrophages following co-culture with *S. aureus* infected or H_2_O_2_-exposed macrophages.** Bar plot showing the top enriched KEGG pathways among genes associated with the differentially methylated CpG sites (top 10 pathways) for macrophages co-cultured with (a) *S. aureus*-infected macrophages. **(b)** H_2_O_2_-exposed macrophages.

**Supplementary figure 5. Macrophage control of *Mycobacterium tuberculosis* growth after co-culture with BCG-infected, *S. aureus-*infected or H_2_O_2_-exposed macrophages.** Experiments were performed on macrophages isolated from blood of four donors. Naïve macrophage was co-cultured with *M. bovis* (BCG)-infected, *S. aureus* (SA)-infected or H_2_O_2_-exposed macrophages during differentiation and subsequently infected with *M. tuberculosis* (H37Rv) expressing GFP. Bacterial growth was measured in a live cell imaging microscope by fluorescent intensity over five days. The values are normalized to day 0.

**Supplementary figure 6. Effect of conditioned media from macrophages under different exposures depleted of extracellular vesicles (EVs) by ultracentrifugation compared to conditioned media containing EVs on the mycobacterial control.** Monocytes from two donors were differentiated in the presence of either; ultracentrifugated EV depleted conditioned media (black line and square) or conditioned media containing EVs (colored line and circle). Condition media was obtained from macrophage cultures infected with *S. aureus* (SA), exposed to H_2_O_2_ or left untreated. After differentiation, the macrophages were subsequently infected with GFP-expressing *M. tuberculosis* (H37Rv). Bacterial burden was assessed by quantifying the fold change in H37Rv-GFP fluorescence over five days.

