## Supplementary Methods for "Epigenetic Crosstalk Between BCG-Infected Macrophages and Naïve Monocytes Potentiates Antimycobacterial Activity"

*) Corresponding author

**Material and Methods**

**Study Design:**

In this study, we investigated how BCG-infected macrophages can modulate the functional behavior of healthy macrophages against Mtb. In addition, we focused on elucidating the mechanisms by which extracellular vesicles (EVs) released by infected macrophages contribute to these modulatory effects. For this purpose, we established a transwell co-culture model. We used Staphylococcus aureus (SA) and Hydrogen peroxide (H₂O₂) in our model as distinct activators of macrophage inflammatory and antimicrobial stimulation.

In this system, differentiated macrophages (MQs) were seeded on transwell inserts (0.4 µm pore size) and stimulated with BCG, SA or H₂O₂. In parallel, naive monocytes were seeded in the lower chamber, where they matured in the presence of released EVs and other factors by the infected MQs. After maturation, these matured recipient MQs were infected with Mtb to assess their antimicrobial activity against Mtb. The bactericidal capacity of recipient MQs was then monitored in real time using live-cell imaging of GFP-expressing Mtb.

To evaluate potential epigenetic reprogramming in the recipient MQs, genomic DNA was extracted and subjected to DNA methylation profiling.

In addition, to characterize the potential signals transferred by the infection-induced EVs, supernatants were collected after MQ stimulation with BCG, SA, or H₂O₂. EVs were isolated and analyzed by nanoparticle tracking analysis (NTA), transmission electron microscopy (TEM), and flow cytometry to determine size distribution, morphology, and surface marker expression. Proteomic profiling was then performed on EVs released by BCG-infected MQs as well as other conditions. The identified peptides were mapped to biological pathways to provide insight into their potential roles in EV-mediated intercellular communication and immune modulation.

**Ethics statement**

The buffy coat preparations from whole blood were obtained from healthy volunteers from Linköping University Hospital Blood Bank.

**Preparation of Bacterial Cultures for Macrophage Infection**

BCG cultures (Danish vaccine strain, Statens Serum Institute (SSI), Denmark)) grown for 3 weeks were used for infection. SA (RN4220) was prepared from overnight cultures, followed by a 2-hour subculture, which was used for the experiment. Before infection, the bacterial cultures were centrifuged in 4000g and washed with PBS/0.05 tween and the pellets were resuspended in antibiotic free media passed through 21g needle and OD was measured, the number of bacteria was calculated by:

BCG : (OD +0.155)/0.161 * 10^7^ = 1.95 * 10^7^ /ml	bacteria

SA: (OD*179)-18.5= 137.2*10^6^ /ml

**Human Monocyte Isolation and Macrophage differentiation:**

Peripheral blood mononuclear cells (PBMCs) were isolated from buffy coats (leukocyte-enriched blood fractions) obtained from healthy volunteers (Linköping University Hospital blood bank, Linköping). Isolation was carried out using density gradient centrifugation with Lymphoprep (Axis-Shield, UK) in SepMate-50 tubes (STEMCELL Technologies, Vancouver, BC, Canada) following the manufacturer’s instructions. The isolated mononuclear cells were seeded in culture dishes containing Dulbecco’s Modified Eagle Medium (DMEM)(Gibco) and allowed to adhere for 1–2 hours, after which non-adherent lymphocytes were gently removed by washing. The adherent cells were then maintained in complete DMEM supplemented with 10% fetal bovine serum (FBS), 2 mM GlutaMAX, 100 U/mL penicillin, and 100 μg/mL streptomycin (all from Gibco) and 50ng/ml MCSF (Invitrogen, Carlsbad, CA, USA) and were allowed to differentiate for 6 days.

**Preparation of recipient macrophages by co-culture models:** On day 6, macrophages were detached using 0.05% trypsin-EDTA (Gibco), counted, and reseeded at 120,000 cells per well into transwell inserts (Corning Inc., Corning, NY, USA) in antibiotic-free EV-free cell culture medium. Macrophages were rested overnight before treatment. One day post seeding, MQs on the inserts were exposed to either BCG or SA at multiplicities of infection (MOI) of 10 or 30 for 2 hours or 30 minutes, respectively. A separate group was treated with 120 µM hydrogen peroxide (H2O2) for 24 hours. Control cells were incubated with Dulbecco’s Phosphate-Buffered Saline (DPBS)(Gibco). After exposure, cells were washed three times with DPBS and kept in fresh complete media containing EV-depleted FBS (Gibco). In parallel, naive monocytes freshly isolated from buffy coats were seeded in the lower chambers of 12-well plates, onto which the transwell inserts were transferred. After 3 days of coculture, the inserts were removed, and the media in the lower chambers was refreshed. These monocytes were subsequently matured into macrophages and infected with Mycobacterium tuberculosis on day 7 post-seeding**.**

**Control Experiments:** For control experiments, media from different treatment conditions were subjected to ultracentrifugation (UC-Optima XPN-90 ultracentrifuge, Beckman Coulter, Brea, CA, USA)) to remove extracellular vesicles (EVs). Naive monocytes were seeded in 96-well plates at a density of 10,000 cells per well and allowed to adhere for 2 hours. Then, the media was replaced with 50% fresh complete media supplemented with 100 ng/ml M-CSF and 20% EV-depleted FBS, combined with 50% of the pre-prepared EV depleted media. After 3 days of culture, the whole media was refreshed with fresh complete media. On day 7 post-isolation, matured macrophages were infected with Mycobacterium tuberculosis.

**Culture and Infection of Macrophages with Mycobacterium tuberculosis H37Rv-GFP**

The laboratory strain *Mycobacterium tuberculosis* H37Rv, carrying the plasmid pFPV2 encoding green fluorescent protein (GFP) (designated as Mtb-GFP), was cultured in Middlebrook 7H9 broth supplemented with albumin-dextrose-catalase (Becton-Dickinson), 0.05% Tween 80, and kanamycin (20 μg/mL) at 37 °C for 2–3 weeks. The culture was then transferred to fresh medium and incubated for an additional 7 days to obtain bacteria in the early logarithmic growth phase. For infection assays, bacterial cells were collected, washed, and resuspended in antibiotic-free complete DMEM. To eliminate clumps, the suspension was passed through a 5.0 µm Millex®-SV syringe filter (Merck). Macrophages were subsequently infected with Mtb at a multiplicity of infection (MOI) of 1.

**Live-Cell Microscopy of** **Mtb-Infected Macrophages**
 Macrophages infected with the Mtb H37Rv-GFP strain were analyzed using the IncuCyte S3 live-cell analysis system (Sartorius, Göttingen, Germany). Fluorescence signals corresponding to bacterial growth were recorded every 6 hours, capturing four images per well at 20× magnification, continuing until 5 days post-infection. Data were processed using the IncuCyte S3 software and quantified as the total integrated intensity of green, fluorescent objects (green calibration units × μm² per image).

**DNA Extraction and DNA Methylation Data Processing:**

**DNA extraction :**

Genomic DNA was isolated from the recipient MQs using the DNA extraction kit (QIAamp DNA mini kit, Qiagen, Hilden, Germany). DNA concentration was measured with the Quantus Fluorometer and QuantiFluor ONE dsDNA System (E4871, Promega, USA), and DNA quality was evaluated using a NanoDrop ND-1000 spectrophotometer (Thermo Fisher Scientific, USA).

**Methylation array data:**

Genomic DNA (250 ng/sample) samples were bisulfite-converted using EZ DNA Methylation Kit (Zymo Research) and profiled using the Illumina MethylationEPIC v1.0 array. BeadChips were scanned on an Illumina NextSeq 550.

**Data preprocessing and quality control:**

Methylation signal intensities were extracted from raw IDAT files using the minfi R package (v2.34.0) [1]. Preprocessing and within-array normalization were performed using the Subset-quantile Within Array Normalization (SWAN) algorithm to reduce technical variability between type I and type II probes and converted to β-values for downstream analysis. [2] )

Probe-level quality control and filtering were applied. Probes identified as cross-reactive, , were excluded. Detection p-values were calculated, and probes with detection p-values greater than 0.01 in any sample were filtered out. Probes overlapping known single nucleotide polymorphisms (SNPs) were filtered out. Genomic mapping and annotation were performed using minfi and IlluminaHumanMethylationEPICanno.ilm10b4.hg19.
Quality control was further assessed using control probe strip plots, QC reports, and density plots of β-value distributions generated in R.

**Differential DNA methylation analysis :**

Differentially methylated CpGs (DMCs) between groups were identified using the limma R package (v2.34.0) [3]. We applied a nominal P value cutoff of 0.05 and a |logFC| cutoff of 0.1 to determine whether a CpG was differentially methylated. The identified DMCs were mapped to corresponding differentially methylated genes (DMGs) using Illumina EPIC array annotations ilm10b4.hg1932. The results were visualized as volcano plots using ggplot2 [4] in R.

**Pathway enrichment analysis and network analysis :**

The genes annotated to identified DMCs were analyzed for pathway enrichment using EnrichR [5] and the Kyoto Encyclopedia of Genes and Genomes (KEGG) database [6]. To enhance visualization and interpretation of enrichment results, a dot plot was generated using the ggplot2 R package [4]. Gene–pathway relationships and network connectivity among enriched pathways were visualized using the clusterProfiler and enrichplot R packages (cnetplot), allowing depiction of shared genes and functional overlaps across pathways.

**EV proteomic and DNAm interactome network analysis**

To investigate potential molecular interactions between extracellular vesicle (EV)–derived proteins and genes associated with differential DNA methylation (DNAm), an integrated interactome network was constructed using the STRINGdb R package (v2.20.0) [7].

A curated dataset containing significantly expressed EV proteins and DMGs with unique identifiers from both datasets were extracted and mapped to the STRING database (version 12.0) for Homo sapiens (taxonomy ID: 9606), using a high-confidence interaction score threshold of 0.7 (700). Unmapped entities were excluded to ensure annotation precision.

Protein–protein interaction (PPI) data were retrieved using the get_interactions() function in STRINGdb. The resulting interaction data were transformed into an undirected network object using igraph (v1.5.1) [8]. The network was visualized with ggraph and ggplot2 (v3.4.0). Node size was scaled by degree centrality, and labels were added using ggrepel to minimize overlap.

Topological characteristics of the EV–DNAm interactome were assessed using igraph functions. Degree centrality was calculated to identify highly connected hub nodes potentially mediating cross-domain molecular communication. Betweenness and closeness centralities were computed to evaluate node influence and network efficiency, respectively.

**Extracellular isolation and characterization:**

**Isolation:** EVs were isolated using either size exclusion chromatography (SEC) with either commercially available IZON qEV columns. Briefly, conditioned media from all experimental groups (H2O2, BCG, SA, and controls) were collected three days post-infection, centrifuged at 2,000 × g for 15 minutes, and filtered twice through 0.22 μm filters( MilliporeSigma, Burlington, MA, USA). The filtered media were then concentrated using Amicon Ultra centrifugal filters (3 kDa molecular weight cutoff )( Millipore) by centrifugation at 4,000 × g for 30 minutes per 15 mL aliquot, yielding approximately 1 mL of concentrated media. This concentrate was loaded onto IZON qEV columns (Izon Science,Christchurch, New Zealand) pre-equilibrated with PBS, and fractions were collected following the manufacturer’s instructions.

**Flowcytometry characterization**: Extracellular vesicle (EV) surface marker profiling was performed using a multiplex bead-based flow cytometry assay (MACSPlex Exosome Kit, Miltenyi Biotec) following the manufacturer’s instructions. Briefly, EV samples were incubated with a panel of fluorescently capture beads, each coated with antibodies specific to 38 exosomal surface markers. After overnight incubation at room temperature, bead-bound EVs were stained with a cocktail of APC-conjugated detection antibodies against CD9, CD63, and CD81.

Sample acquisition was performed on a flow cytometer (Beckman Coulter, Brea, CA, USA). For analysis, individual bead populations were identified based on their fluorescence in the FITC and PE channels, the fluorescence intensity was quantified using FlowJo software version 10 (BD Life Sciences, Ashland, OR) .

**NTA measurement with Nanosight NS300:** Nanoparticle tracking analysis was performed using a NanoSight NS300 instrument (Malvern Panalytical) equipped with a 488 nm, 45 mW laser and sCMOS camera at a constant temperature (23°C) following the manufacturer’s guidelines (Malvern Panalytical. NanoSight NS300 User Manual (version MAN0516-08-EN-00). Malvern Panalytical Ltd. 2016). EV suspensions were diluted in sterile, filtered PBS to achieve an optimal particle concentration for the measurement by the machine according the NanoSight NS300 Software Guide (~10^7–10^9 particles/mL, approximately 20–100 particles in the field of view). For each measurement, 1 mL of diluted sample was loaded into the analysis chamber. All instrument settings, including camera level and detection threshold, were established by pre-testing and then kept constant across all samples. The camera level was adjusted to level 16 to optimize particle visibility without signal saturation, and autofocus settings were used to eliminate indistinct particles. Detection threshold was set on 5 to balance sensitivity and minimize false positive counts, with limits on red and blue particle markers as internal quality control. For each sample, five videos of 60 second duration were recorded at 23°C with a syringe flow rate of 30 µL/s and were processed with NanoSight Software (version 3.4).

**Characterization with TEM:** EV samples were fixed with 1%–2% paraformaldehyde and visualized using negative staining for transmission electron microscopy (TEM) at the Linköping University Core Facility. In brief, 5 μL of each sample was placed on hydrophilic formvar/carbon-coated 300-mesh copper grids (TED PELLA, Inc.). The grids were then washed, blotted, and stained with 2% uranyl acetate. Electron micrographs were captured using a JEOL JEM 1400 Flash transmission electron microscope (80 kV; JEOL Ltd., Tokyo, Japan).

**Proteomics:**

**Proteomics Sample Preparation:** Protein extraction from extracellular vehicles (EVs) was performed using pre-chilled 1× RIPA buffer (Sigma) supplemented with protease inhibitors (Sigma), followed by sonication and centrifugation. Protein concentration was determined by BCA assay.

We used ultrasonic-based filter aided ample preparation (FASP) [9] method for sample preparation for proteomics. 10ug proteins were reduced by adding 200 μL of 10 mM dithiothreitol (DTT) (Sigma), and incubated for 5 min at 95 °C., after chilling at RT, samples were loaded in a 10K Amicon filters (Millipore Sigma), with a 3000 molecular weight cutoff (MWCO). The peptides present in the membrane were washed with 200 μL of 8 M urea (Sigma). Afterward, centrifugation for 20 min at 14 000g was done, followed by protein alkylation by the addition of 100 μL of 50 mM iodoacetamide (IAA)((Sigma)) in 8 M urea and 25 mM AmBic solution (Sigma). The alkylation step was sped up using the ultrasonic microplate horn assembly during 5.25 min (7 cycles: 30 s onand 15 s off UT, 25% UA, 20 kHz UF). Finally, 100 μL of 1:30 trypsin (Sigma) in 12.5 mM AmBic solution was added, and the protein digestion was processed using the ultrasonic for 5.25 min (7 cycles: 30s on and 15 s off UT, 25% UA, 20 kHz UF). Peptides were desalted using C18 tips (Thermo Fisher Scientific, Waltham, MA, USA) and dried by vacuum centrifugation for further use in LC-MS/MS analysis.

**Determination of peptide concentration and quality:** Briefly samples were solved in formic acid(FA) (0.1%) (Sigma) and sonicated with water bath sonicator for 10min and spin 13000g, 10min at RT. Peptide concentration and quality was measured using a nanodrop in A280. Peptide were diluted to a final concentration 0.1ug/ul for LC-MS/Ms analysis.

**LC–MS/MS Analysis:** LC-MS/MS analysis was performed using a High-Resolution Trapped Ion Mobility timsTOF HT mass spectrometer coupled to a nanoElute 2 LC system via a CaptiveSpray ionization source (Bruker Daltonics). The prepared peptid mixture reconstituted in 0.1 % FA , was loaded of reverse-phase C18 HPLC column (PepSep XTREME with dimensions of 25 cm x 150 µm x 1.5 µm, Bruker Daltonics) and peptides were separated in 45 min with a gradient of 0.1% formic acid in water (A) and 0.1% formic acid in acetonitrile (B) as follows: from 2% B to 17% B over 25 minutes; from 25 % B to 37% B over 35minutes; from 37 % B to 95% B over 45 minutes a flow rate of 400 nl/min and a column temperature of 50°C. All data were acquired under the dia-PASEF mode with a MS1 scan range of 100-1700 m/z, and the collision energy was linearly interpolated between 1/K0 values, from 20 eV at 0.6 Vs/cm2 to 59 eV at 1.6 Vs/cm2, keeping constant above or below.

**Protein Identification and Data processing:**

dia-PASEF data files were analyzed in Spectronaut (v 18 Biognosys) [10] using a custom database to identify the peptides in different groups of treatment. The following parameters were applied for the search: trypsin as the proteolytic enzyme, allowing up to one missed cleavage; carbamidomethylation of cysteine as a fixed modification; oxidation of methionine as a variable modification. False discovery rate (FDR) cut-off of 1% was used for peptide and protein level identifications.

Proteomics data in TSV format were imported into R using the *readr* package. Protein presence across four sample groups (BCG, SA, H2O2 and Control) was assessed by counting non-missing, positive abundance values per sample. Proteins uniquely detected in each group were identified and their corresponding gene symbols were extracted. Venn diagrams visualizing protein overlaps between groups were generated with the VennDiagram package [11]. Gene overlaps including pairwise and three-group intersections were calculated using the GeneOverlap package.

For functional interpretation, gene sets unique to each group were subjected to KEGG pathway enrichment analysis using clusterProfiler [12] with gene symbol to Entrez ID conversion based on org.Hs.eg.db. Resulting enrichment data were filtered, summarized, and visualized using ggplot and enrichplot, including cnetplots to display gene-pathway relationships. Bar plots depicting unique gene counts and total detected peptides per group were generated to illustrate protein distribution patterns.

**Statistics and Bioinformatics:**

Statistical analyses on Mtb growth experiments were performed with GraphPad Prism (version 10.5.0) using wilcoxon signed- rank test.

We used linear modeling followed by empirical Bayes moderation, as implemented in the limma package for differential analysis of methylation data.

P-values were adjusted for multiple testing using the Benjamini-Hochberg procedure for False Discovery Rate (FDR) correction at 5%. In case no significant FDR was reached, we used nominal p-value < 0.05.

**Data/code Availability:**

https://github.com/Lerm-Lab/Epigenetic-Crosstalk
