## Supplementary figures and images for "Epigenetic Crosstalk Between BCG-Infected Macrophages and Naïve Monocytes Potentiates Antimycobacterial Activity"

### Supplementary Figure 1

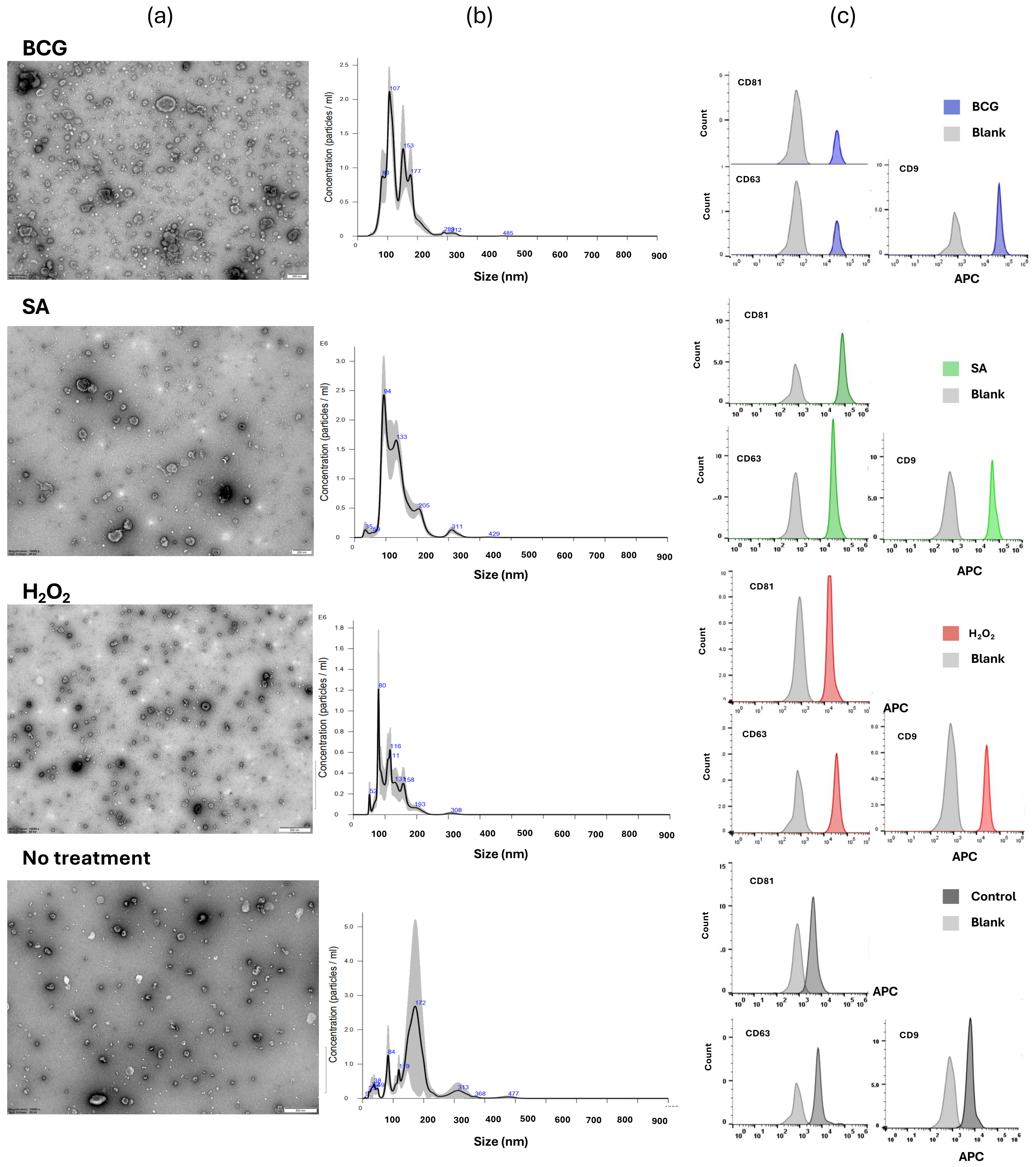


Supplementary figure 1.

### Supplementary Figure 2

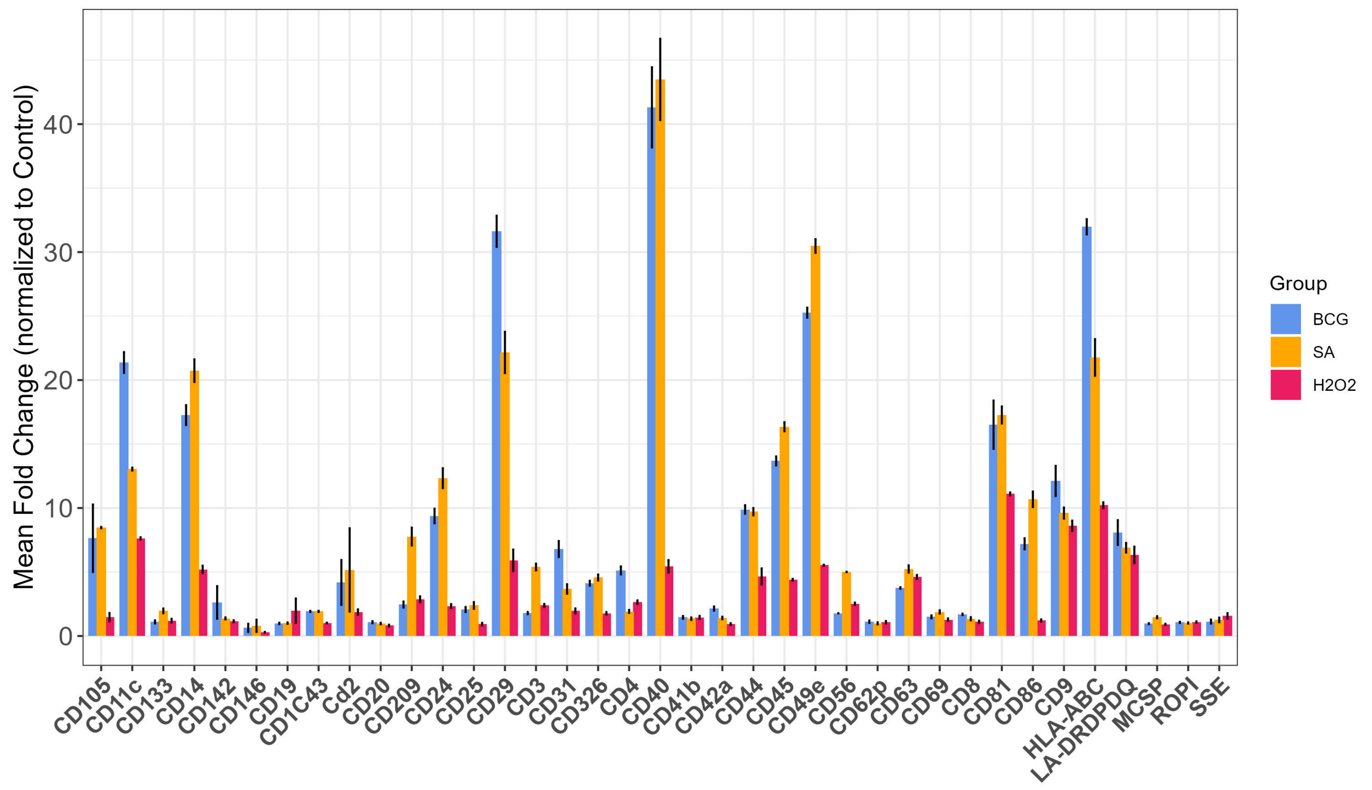


Supplementary figure 2.

### Supplementary Figure 3

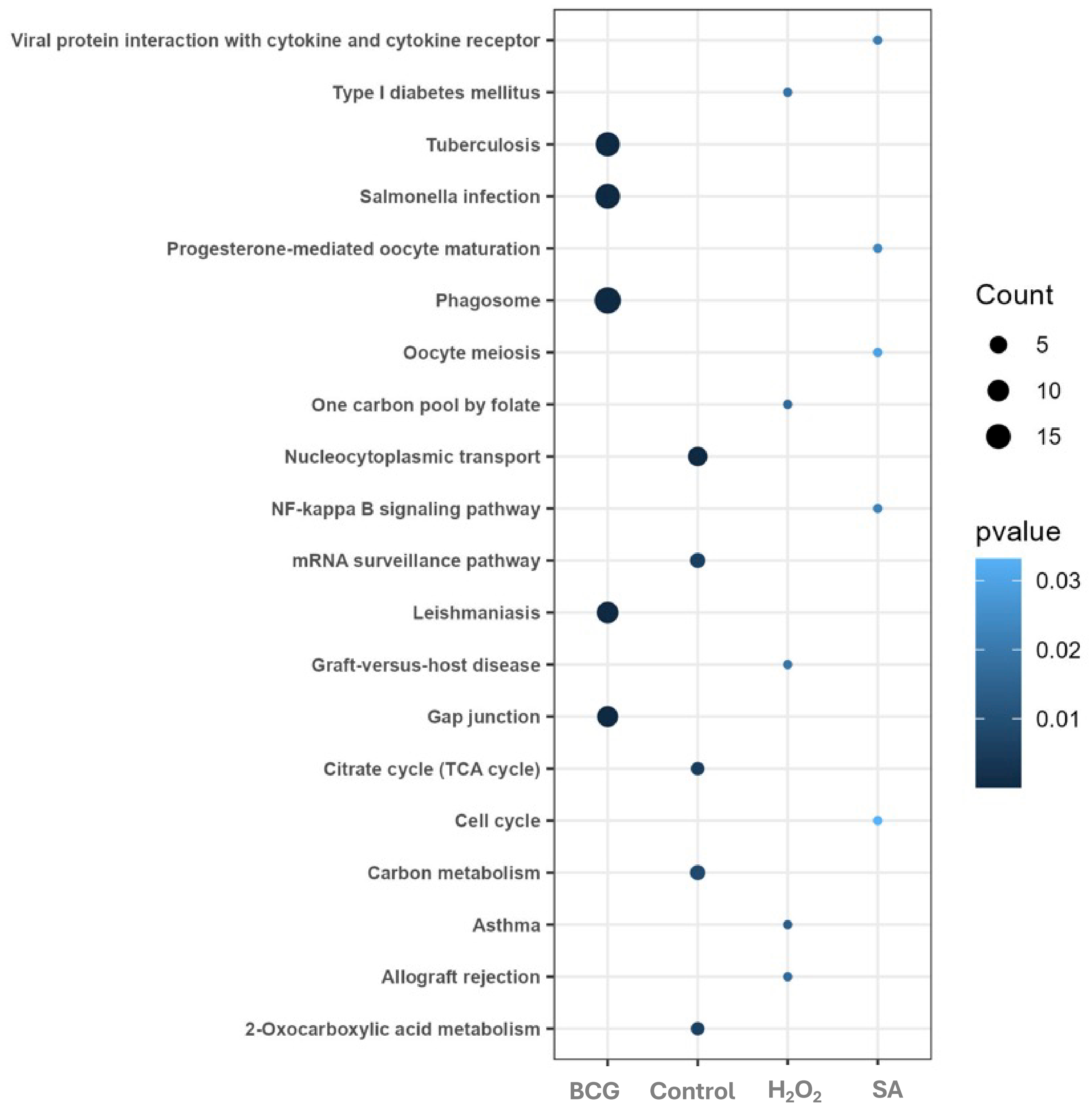


Supplementary figure 3.

### Supplementary Figure 4

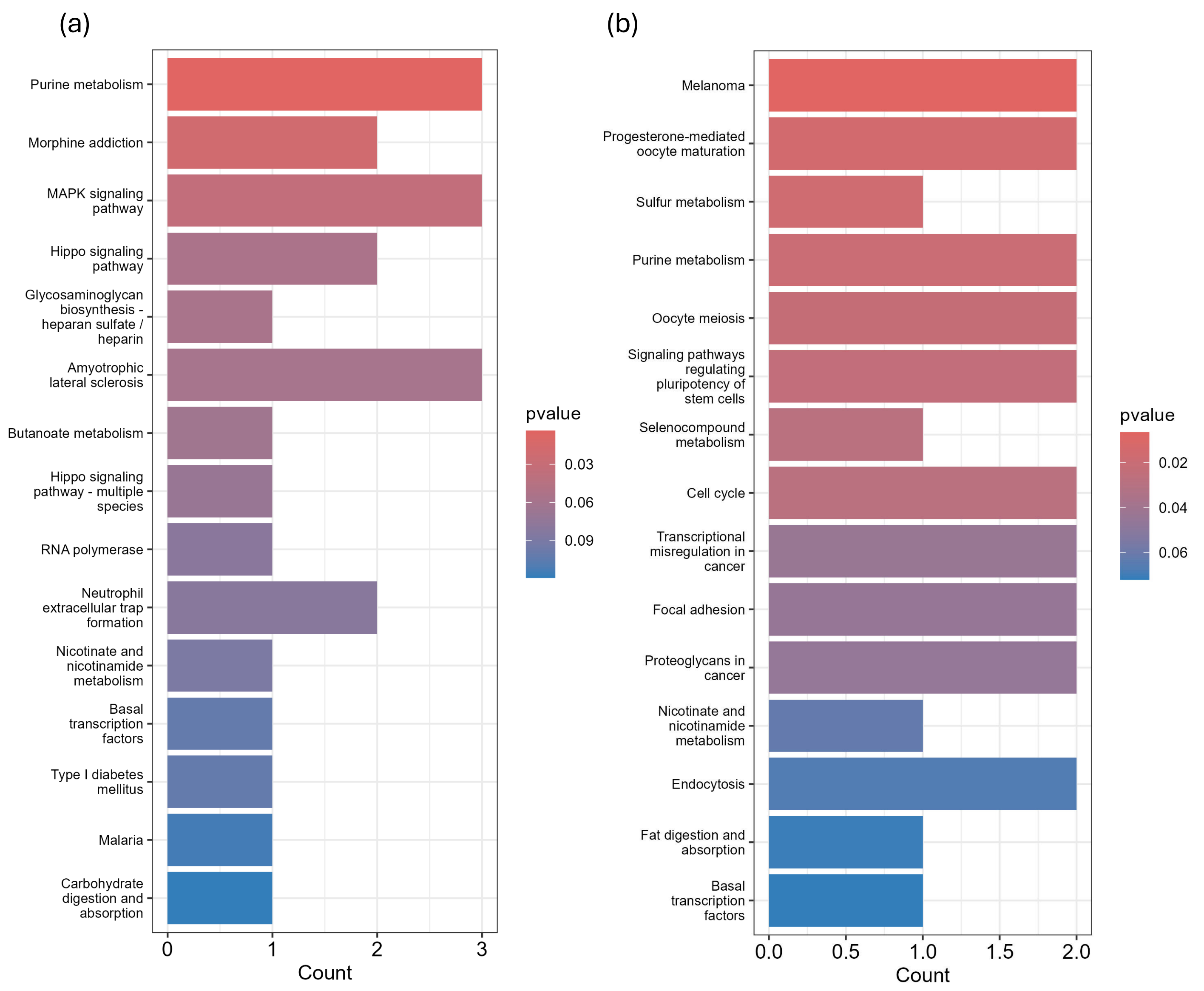


Supplementary figure 4.

### Supplementary Figure 5

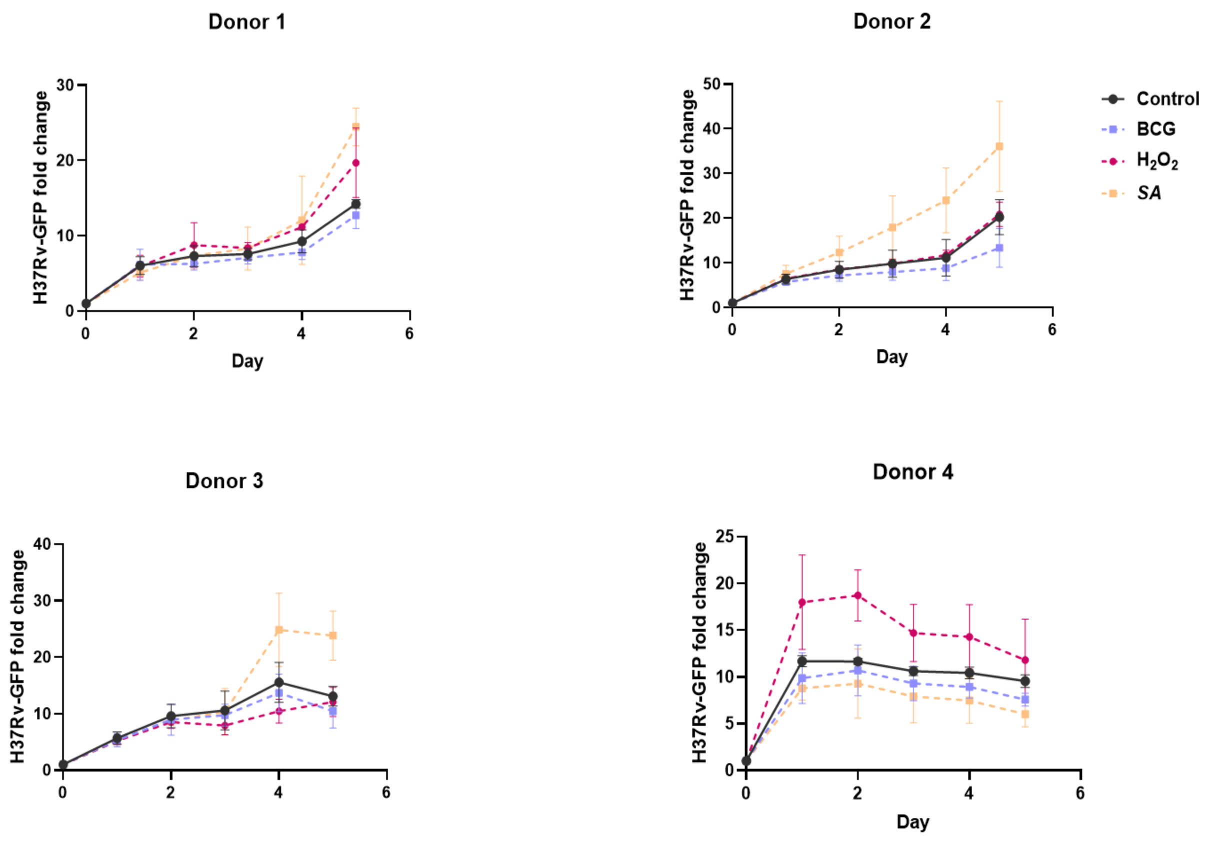


Supplementary figure 5.

### Supplementary Figure 6

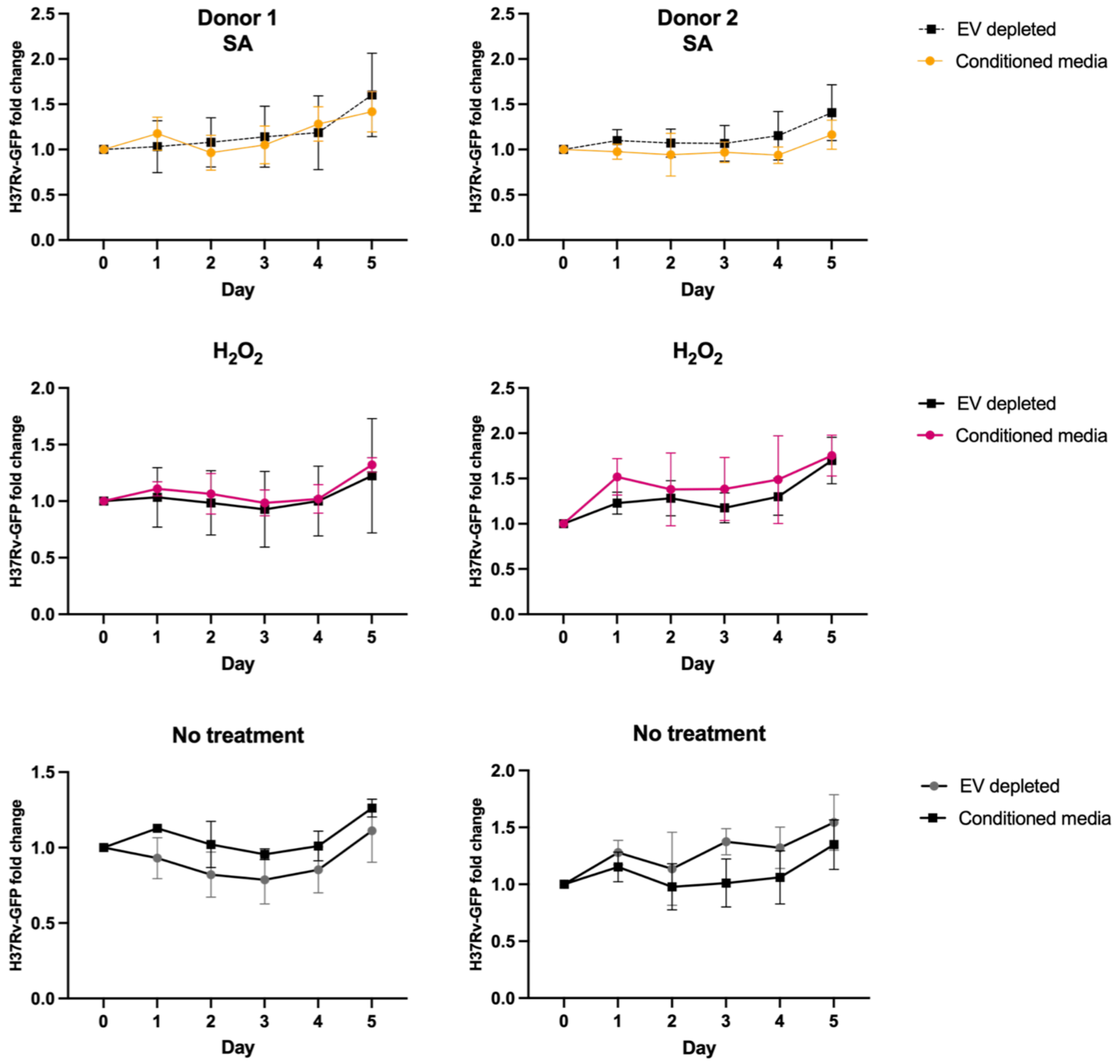


Supplementary figure 6.
